# Transient Magnetothermal Neuronal Silencing using the Chloride Channel Anoctamin1 (TMEM16A)

**DOI:** 10.1101/306555

**Authors:** Rahul Munshi, Shahnaz Qadri, Arnd Pralle

## Abstract

The importance of specific neurons to a network’s function is best studied by precisely timed, reversible silencing of these neurons. Previously, we showed that alternating magnetic field mediated heating of magnetic nanoparticles bound to neurons expressing temperature-sensitive cation channels TRPV1, stimulates these neurons to fire and affects animal behavior in vivo (Munshi et al., 2017). Here, we demonstrate how to apply magnetic nanoparticle mediated heating to silence target neurons. Rat hippocampal neuron cultures are transfected to express the temperature gated chloride Anoctamin1 (TMEM16A) channels. Within seconds the heating of the membrane opens the Anoctamin1 (TMEM16A) channels, suppressing action potential firing. Five seconds of magnetic field application leads to about 12 seconds of silencing, with a latency of about 2 seconds and an average suppression ratio of more than 80%. The method provides a promising avenue for tether free, remote, transient neuronal silencing in vivo for both scientific and therapeutic applications.

## Introduction

Spatially and temporally tightly regulated signaling in complex brain circuits control behavior, emotions and all brain function. Tools to modulate specific network components and connections are absolutely crucial in the study of the functional connectivity of these circuit. In the past decade, electrical, optical, chemical, and magnetic tools to activate specific neurons deep in the brain have been developed (Deisseroth, 2015; Kim, Adhikari, & Deisseroth, 2017; Warden, Cardin, & Deisseroth, 2014). However, the only sure test whether a network component is essential, is to transiently silence that component (Nomura et al., 2015). So far, only a few silencing approaches have been developed (Chow et al., 2010; Okazaki, Sudo, & Takagi, 2012; Slimko, Mckinney, Anderson, Davidson, & Lester, 2002). Silencing is typically achieved by maintaining the membrane potential negative enough to suppress action potential firing (Hodge, 2009). Pharmacological approaches use engineered small molecules to manipulate G-protein signaling pathways (Stachniak, Ghosh, & Sternson, 2014) or ligand gated ion channels to subdue action potentials (Lerchner et al., 2007; Slimko et al., 2002). Optogenetic tools employ modified opsins as light driven inward chloride pumps or outward proton pumps (Chow et al., 2010; Okazaki, Takahashi, Toyoda, & Takagi, 2014; Raimondo, Kay, Ellender, & Akerman, 2012; Sudo et al., 2013). More recently, a chloride conducting channelrhodopsin was developed (E. G. Govorunova, Sineshchekov, Janz, Liu, & Spudich, 2015; Elena G. Govorunova et al., 2017; J. Wietek et al., 2014; Jonas Wietek et al., 2015), which provided a light gated ion channel based silencing mechanism.

Photothermally induced hyperpolarization in wild type neurons have also been achieved by interfacing with a variety of engineered materials, ranging from conjugated polymers (Feyen et al., 2016) to plasmonic gold nanorods (Yoo, Hong, Choi, Park, & Nam, 2014). Upconversion nanoparticles, which absorb in the near infrared region and emit in the wavelength range of inhibitory opsins, has been used to remotely silence deep brain neurons (X. Lin et al., 2018).

Here, we introduce magnetothermal silencing, using the heat generated by superparamagnetic nanoparticles to activate the thermosensitive chloride channel Anoctamin 1 (TMEM16A) (Cho et al., 2012; Yang et al., 2008). We co-transfected membrane protein Ano1/TMEM16A and cytosolic calcium indicating protein GCaMP6f (Chen et al., 2013) in rat hippocampal cultures. Polymer encapsuled superparamagnetic nanoparticles were bound to the neuronal plasma membrane via A2B5 antibodies. When exposed to alternating magnetic fields, these particles get heated, raising the membrane temperature (Rosensweig, 2002). The elevated membrane temperatures opened Ano1/TMEM16A channels causing inhibition of calcium spiking recorded by the GCaMP6f signal. Within seconds of turning the magnetic field off, the membrane cooled off and calcium spiking resumed. As Ano1/TMEM16A is a chloride channel naturally occurring in mammals (Cho et al., 2012; F. Huang et al., 2009; Terashima, Picollo, & Accardi, 2013), no protein engineering or chemical manipulation is needed, as opposed to the use of microbial opsins (Berndt, Lee, Ramakrishnan, & Deissoroth, 2014; Hodge, 2009; Jonas Wietek et al., 2015). This method can be used for deep brain silencing, as it uses the same alternating magnetic fields, nanoparticles and nanoparticle heating as our prior report on stimulation in vivo (Munshi et al., 2017). Only cells expressing Ano1/TMEM16A channels and having membrane bound nanoparticles show inactivation, while the adjoining cells remain unaffected. Thus, it can serve as a truly remote silencing technique with neuronal subpopulation level cell selectivity.

## Results

### Thermal suppression of Ca^2+^influx in Ano1/TMEM16A^+^neurons

We first determined whether spontaneously firing Ano1/TMEM16A^+^ neurons could be significantly inactivated at higher temperatures. To demonstrate thermal suppression of intracellular Ca^2+^ influx, we co-transfected (see Methods) rat hippocampal neurons with plasmid DNAs for mAno1/TMEM16A-mCherry and the genetic calcium indicator GCaMP6f. Experiments were performed 48 hours after transfection. The cultured neurons were treated with 1μM Tetrodotoxin (TTX, Sigma-Aldrich), 24 hours prior to experiments. Right before the experiments, the TTX was washed out with high K^+^ imaging buffer (NaCl 145, CaCl_2_ 2, MgCl_2_ 1, KCl 4.5, HEPES 10, Glucose 20 (all in mM)). Experiments were started only after regular Ca^2+^ activity was observed, following the TTX wash. Cells expressing Ano1/TMEM16A were identified by the mCherry marker (Figure 1A, Figure 1 Figure supplement 1, 2A). GCaMP6f fluorescence signal captures cytosolic Calcium transients, resulting from membrane depolarization (Chen et al., 2013). The average GCaMP6f peak, corresponding to an isolated action potential (AP) firing was indistinguishable between neurons with and without Ano1/TMEM16A. Measured at 37 °C, the rise times (duration until calcium peak is reached) were 0.275 ± 0.048 s and 0.250 ± 0.029 s (p = 0.674), and the half-decay time for the Ca^2+^ decay were 0.274 ± 0.011 s and 0.290 ± 0.055 s (p = 0.801) in Ano1/TMEM16A^+/-^ neurons respectively (Figure 1 – Figure Supplement 2 B, C).

**Figure 1.**
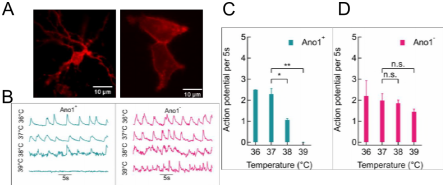
Thermal silencing of Anoctamin 1^+^ hippocampal neurons. **(A)** Anol expression in (i) rat hippocampal neurons and (ii) HEK293 cells, visualized using mCherry tag.**(B)** Comparison of representative GCaMP6f traces between Ano1^+/-^ neurons at 36, 37, 38 and 39 °C. **(C, D)** Action potential firing rate (per 5s) comparison in Ano1 ^+/-^ neurons at various bath temperatures. Significant suppression in firing rate was observed at 38 °C (p = 0.0163) and 39 °C (p = 0.0032) in Ano1^+^ neurons, compared to 37 °C (T-test). In Ano1^-^ neurons similar comparison yielded nonsignificant changes (p = 0.7067 and p = 0.1682 at 38 °C and 39 °C respectively).

Representative GCaMP6f fluorescence intensity traces recorded from cells with and without Ano1/TMEM16A at various steady bath temperatures (36°, 37°, 38° and 39°C) are shown in Figure 1B. A sharp decline in GCaMP6f peaks was seen in Ano1/TMEM16A^+^ traces at 38° C and more significantly at 39 °C, where the peaks completely disappeared. On the other hand, in cells not expressing Ano1/TMEM16A, no such suppression in Ca^2+^ activities was observed. To quantify thermal suppression, we calculated AP firing rates from GCaMP6f fluorescence intensity traces by deconvolving the calcium transients (see Methods, and Figure 3 – Figure Supplement 1). The five second AP firing rates in Ano1/TMEM16A^+^ cells were significantly reduced from 37 °C to 39 °C: from 2.29 ± 0.26 at 37 °C to 1.06 ± 0.06 at 38 °C and 0 at 39 °C (p = 0.016 at 38 °C (n = 4) and p = 0.003 at 39 °C (n = 5) compared to 37 °C, unpaired T-test) (Figure 1C). On the other hand, the AP firing rate of Ano1/TMEM16A^-^ neurons did not change over the same temperature range (AP rates averaged over five seconds (n = 5) were 2.20 ± 0.72 at 36 °C, 1.98 ± 0.32 at 37°C, 1.85 ± 0.15 at 38°C and 1.45 ± 0.13 at 39°C). The p-values of rates, compared to 37°C was 0.707 at 38°C and 0.168 at 39°C, obtained by unpaired T-test (Figure 1D).

The firing rates of neurons with Ano1/TMEM16A were indistinguishable from the rates of neurons without the channel at 36 °C and 37 °C (p = 0.663 and 0.453, respectively). However, at at 38 °C and 39 °C they deviated significantly (p = 0.0227 and 0.0051, respectively) (Figure 1 – Figure Supplement 2D). Furthermore, Ano1/TMEM16A^-^ cells fired at 40 °C at 1.38 ± 0.61 APs per five seconds. The extracellular calcium concentration was fixed at 2mM for all the experiments. This establishes that AP firing in Ano1/TMEM16A overexpressing neurons is significantly suppressed at temperatures above 37 °C, while wild type neurons show no such change.

### Heating Neuronal Plasma Membrane with Nanoparticles

For remote magnetothermal silencing of Ano1/TMEM16A^+^ neurons, we applied superparamagnetic nanoparticles to the cell membrane as local heating agents. Core-shell nanoparticles with a 6.7 ± 1.0 nm MnFe_2_O_4_ core, a CoFe_2_O_4_ shell and a total inorganic diameter of 12.9 ± 1.4 nm were encapsulated in PMA (dodecyl-*grafted*-poly-(isobutylene-*alt*-maleic-anhydride)(C. A. J. Lin et al., 2008; Zhang et al., 2015). The specific loss in power (SLP) of these particles, suspended in water, was 553 ± 10 W/g in a 37 kA/m alternating magnetic field (AMF), driven at 412.5 kHz. The outer PMA layer was functionalized with Neutravidin labeled with Alexa Fluor 647 fluorophore (see Methods). To specifically target nanoparticles to the neuronal cell membrane, we briefly incubated neurons first with biotinylated A2B5 antibodies. Then the Neutravidin-dye tagged nanoparticles were added (see Methods). After washing, only the membrane bound nanoparticles remained (Figure 2A), effectively creating a shape conforming array of nanoscopic heat sources on the cell membrane (H. Huang, Delikanli, Zeng, Ferkey, & Pralle, 2010; Munshi et al., 2017). Under alternating magnetic fields, these nanoparticles heated the membrane, opening Ano1/TMEM16A channels (principle, Figure 2B). To measure the temperature change of the magnetothermally-heated cell membrane, we employed autofocus stabilized epifluorescence microscopy under alternating magnetic fields. Time series imaging of the fluorescent tag (Alexa Fluor 647) of the membrane bound nanoparticles (Figure 2 – Figure Supplement 1) enabled measuring temperature changes close to the membrane. In the temperature range of interest, the fluorescence emission intensity of the Alexa Fluor 647 fluorophore decreases approximately linearly with increasing temperature. The rate of normalized intensity change versus temperature was −0.46% °C^−1^. Five seconds of AMF application (300 ± 13 Gauss, 412.5 kHz) was sufficient to raise the temperature by 3.42 ± 0.17 °C (Figure 2C), corresponding to a heating rate of 0.75 ± 0.03 °C/s. Cooling of the membrane starts instantaneously with the turning off of the AMF. With an offset of two seconds from the start of the AMF, we binned time into 5 seconds intervals. Temperature was 38.5 – 40 °C (during 2 – 7 s), 38 – 40 °C (during 7 – 12 s), and 37 – 38 °C (during 17 – 22 s). These time points are marked on the top axis of Figure 2D, for visual guidance. It is to be noted here that the AMF application lasts from 0 – 5 s, and the cooling of the membrane starts instantaneously with the turning off of the AMF.

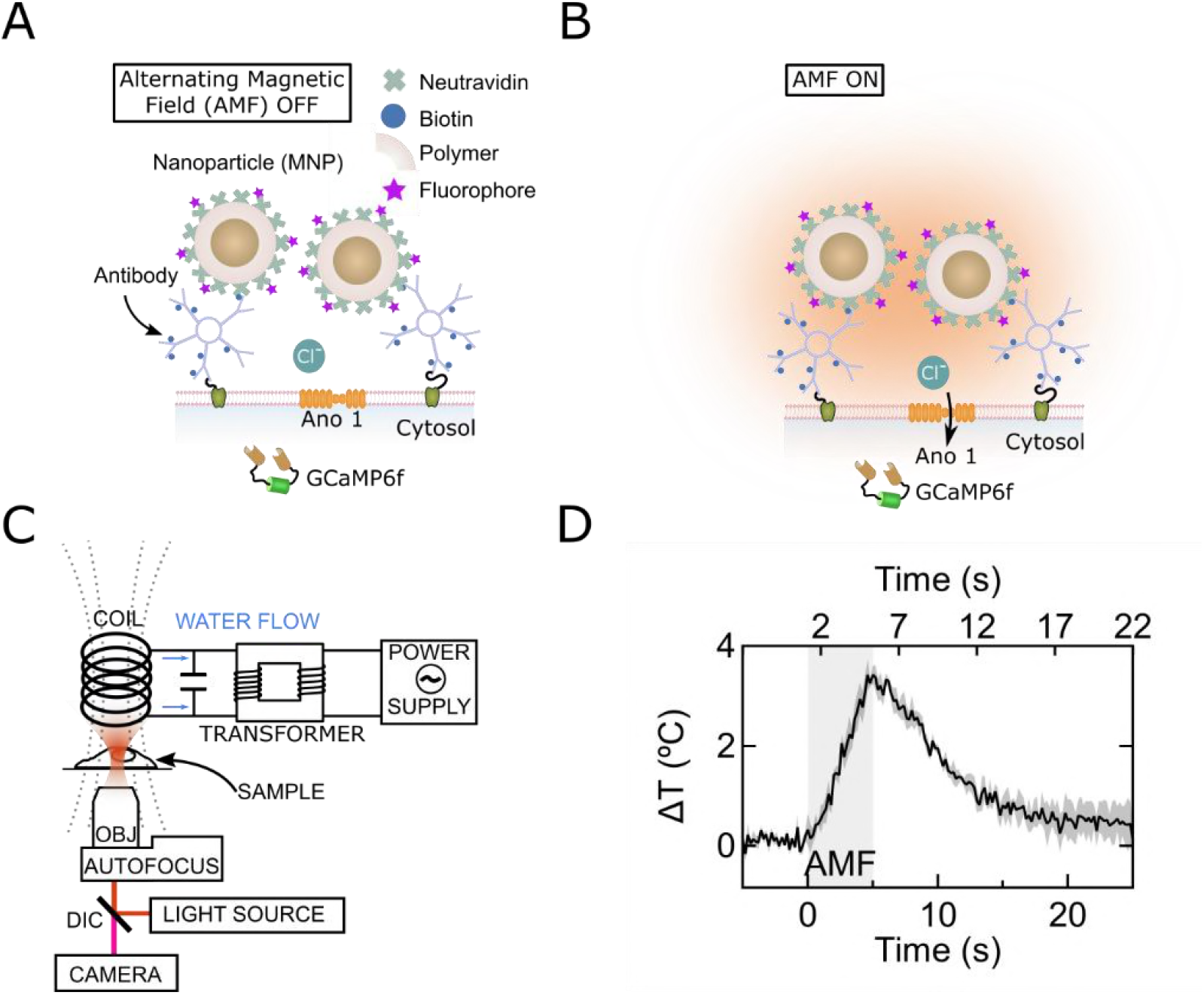
Magneto-thermal silencing scheme. **(A)** and **(B)** illustrate the principle of magnetogenetic silencing. Magnetic nanoparticles (brown) were encapsulated in polymer (PMMA). Fluorescent dye (Alexa Fluor 647) molecules were bound to Neutravidin molecules (green; 5:1 molar ratio). The Neutravidin molecules were covalently attached to the nanoparticle polymer coating. These particles were then attached to the neuronal cell membranes via biotin-avidin bonding formed with the membrane attached biotinylated IgM antibodies. When exposed to alternating magnetic fields (AMF), the particles heat, opening the temperature gated membrane Ano1 channels (B). This causes an influx of Chloride ions, leading to membrane hyperpolarization. **(C)** Schematic, showing principle of magnetic field generation and microscopy under alternating magnetic fields. The field generating coil is placed over the imaging chamber. Autofocus is used to stabilize the objective focus, during magnetic field application. **(D)** Plot shows average temperature rise in magnetic nanoparticle decorated membrane under five seconds magnetic field (n = 3) Bottom axis shows time relative to AMF start, and top axis shows five second time bins when significant temperature points are achieved (e.g. 2−7s = 1- 3 °C, 7−12s = 3- 1 °C, 17- 22s <1°C).

### Magnetothermal Silencing of Ano1/TMEM16A^+^ neurons

To demonstrate transient magnetothermal, Ano1/TMEM16A mediated silencing of neuronal activity we co-transfected cultured, mature rat hippocampal neurons with mAno1/TMEM16A-mcherry and GCaMP6f plasmid DNAs, 48 hours prior to the experiment. 1 μM TTX was added to the culture medium 24 hours prior to the experiment (see Methods). Before the experiment, Alexa Fluor 647 labeled magnetic nanoparticles (MNPs) were bound to the neuronal membrane, using biotinylated A2B5 antibodies (see Methods). Ano1/TMEM16A^+^ neurons were identified via mCherry marker. Cells were washed with high K^+^ imaging buffer to remove any remaining TTX molecules as well as any unbound antibodies or MNPs. Spontaneously firing neurons were selected by their GCaMP6f signal.

Perfusion of the high K^+^ imaging buffer was maintained at 2 ml/minute and the temperature of the sample holder was held at 37 °C by a temperature controlled inline heater. AMF (300 ± 13 Gauss, 412.5 kHz) was applied for 5 seconds intervals using a magnetic hyperthermia coil placed over the Delrin sample holder. Autofocus stabilized GCaMP6f fluorescence images were acquired by a camera at a sampling rate of 10 Hz (schematic, Figure 2C). Average GCaMP6f fluorescence traces, obtained as mean intensity of the pixels spanning the soma of neurons clearly showed suppression of Ca^2+^ influx upon AMF application in Ano1/TMEM16A^+^ neurons (representative traces, Figure 3A). Computed AP events (shown with black bars, under each GCaMP6f fluorescence trace, Figure 3A) show complete silencing in most cases. The AP events were pooled from Ano1/TMEM16A^+^, MNP^+^ neurons from different cultures (n = 9), where cells with different initial spiking rates (at 37 °C) were chosen. Firing rates (mean ± sem, Hz), binned over two second intervals are shown in Figure 3B. Similar graphs for Ano1/TMEM16A^-^, MNP^+^ (n = 15) and Ano1/TMEM16A^+^, MNP^-^ (n = 5) neurons are shown in Figures 3C and D, respectively).

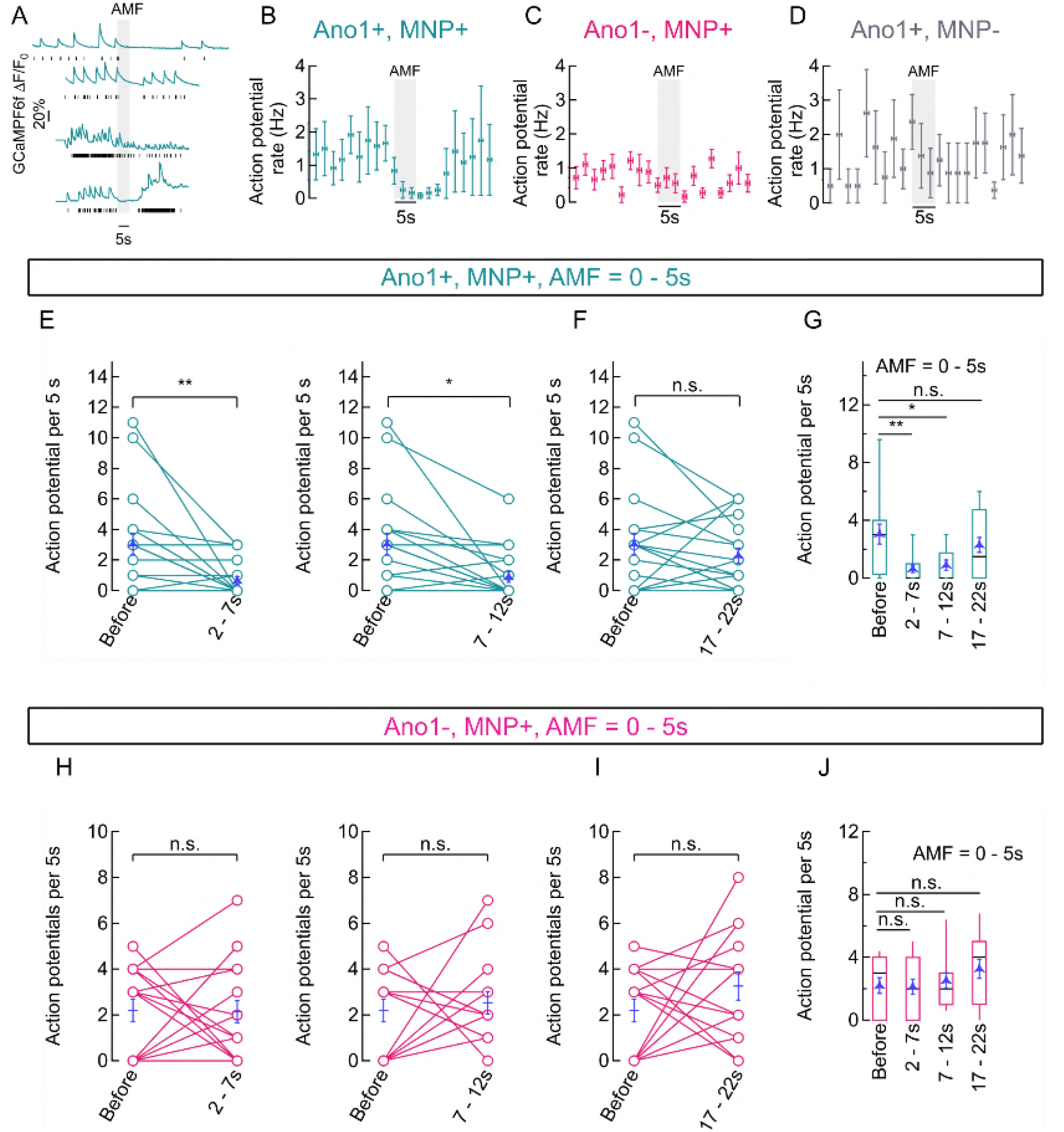
Magneto-thermal silencing of hippocampal neurons. **(A)** Representative traces of GCaMP6f fluorescence intensity averaged over pixels containing the soma of Ano1/TMEM16A+, MNP+ neurons, exposed to AMF (300 ± 13 Gauss, 412.5 kHz; gray, solid). Black sticks under each plot indicate respective calculated action potential (AP) events. (B) Binned (mean ± sem, n = 9) AP firing rates (binned over two seconds, Hz) of Ano1/TMEM16A+, MNP-; neurons. Starting temperature was 37 °C and AMF (300 ± 13 Gauss, 412.5 kHz; gray, solid) was applied for five seconds. **(C, D)** Binned (mean ± sem, n = 9) AP firing rates (2s bins, Hz) of Ano1/TMEM16A-, MNP+ (n = 15, **C**) and Ano1/TMEM16A+, MNP-(n = 5, **D**) neurons are shown. Starting temperature was 37 °C and AMF (300 ± 13 Gauss, 412.5 kHz; gray, solid) was applied for five seconds in both cases. **(E, F)** Shows the comparison between AP firing rates recorded in 5s time intervals corresponding to various temperature ranges achieved during Magnetothermal membrane heating (and subsequent cooling) in Ano1/TMEM16A+, MNP+ neurons (see Figure 2D). Connected plots in **E** compares AP firing rates at different time bins after AMF application (starting at 0 s) with AP rates before AMF, in Ano1/TMEM16A+, MNP+ neurons. Overlaid blue markers show mean ± sem (n = 20). AP rates *before* AMF application (37 °C) was 3.05 ± 0.68 AP per 5s; AP rates during *2- 7s* (38- 40 °C) was 0.65± 0.25 AP per 5s; AP rates during *7- 12s* (38- 39 °C) was 0.90 ± 0.35 AP per 5s; and AP rates during *17- 22s* (37- 38 °C) was 2.30 ± 0.51 AP per 5s. Significance level of each comparison is indicated alongside. The firing rate suppression was significant in *2- 7s* and *7- 12s* intervals (p = 0.0032 and 0.0094 respectively), while insignificant rate change was observed in the *17−22s* bin (p = 0.3858), unpaired T-test. Box and whisker plot in **F** summarizes the results in **E**. Boxes span from 2^nd^ to 3^rd^ quartile (box top, 75% and box bottom, 25%), while whiskers indicate 10^th^ and 90^th^ percentiles. The black lines dividing the boxes indicate the median, while mean ± sem values are overlaid in blue. **(G, H)** Shows the comparison between AP firing rates recorded in 5s time intervals corresponding to various temperatureranges achieved during Magnetothermal membrane heating (and subsequent cooling) in Ano1/TMEM16A-, MNP+ neurons. Connected plots in **G** compares AP firing rates at different time bins after AMF application (starting at 0 s) with AP rates before AMF, in Ano1/TMEM16A-, MNP+ neurons. Overlaid blue markers show mean ± sem (n = 15). AP rates *before* AMF application (37 °C) was 2.20 ± 0.50 AP per 5s; AP rates during *2- 7s* (38- 40 °C) was 2.13 ± 0.48 AP per 5s; AP rates during *7- 12s* (38- 39 °C) was 2.53 ± 0.49 AP per 5s; and AP rates during *17- 22s* (37- 38 °C) was 3.53 ± 0.61 AP per 5s. Significance level of each comparison is indicated alongside. The firing rate change was significant in all cases: *2- 7s, p = 0.9245; 7- 12s*, p = 0.6364; and *17- 22s*, p = 0.1044, unpaired T-test. Box and whisker plot in **H** summarizes the results in **G**. Boxes span from 2^nd^ to 3^rd^ quartile (box top, 75% and box bottom, 25%), while whiskers indicate 10^th^ and 90^th^ percentiles. The black lines dividing the boxes indicate the median, while mean ± sem values are overlaid in blue.

To quantify the silencing, visibly achieved only in Ano1/TMEM16A^+^, MNP^+^ neurons, we compared the computed AP firing rates (Figure 3 – Figure Supplement 1) within time-bins (AMF = 300 ± 13 Gauss, 412.5 kHz; 0- 5s) corresponding to different temperature ranges (as mentioned in the previous section). Firing rates, recorded in the 2- 7 s interval (0.65 ± 0.25 AP per 5s; temperature = 38- 40 °C), were significantly different from the base value (3.05 ± 0.68 AP per 5s; temperature 37 °C), with p = 0.0032. Similarly, firing rates, recorded in the 7- 12 s interval (0.90 ± 0.35 AP per 5s; temperature = 38- 39 °C), were significantly different from the base value (3.05 ± 0.68 AP per 5s; temperature 37 °C), with p = 0.0094 at a suppression ratio, *S = 70.5%*. Firing was observed to resume in the 17- 22 s interval (2.30 ± 0.51 AP per 5s; temperature = 37- 38 °C), with a p-value of 0.3858. All p-values were calculated by unpaired T-test and n = 20 in all cases (Figure 3E, F).

In contrast, Ano1/TMEM16A^-^, MNP^+^ neurons showed no significant change in firing rates with AMF heating. Firing rate at 37 °C, before AMF application was 2.20 ± 0.50 AP per 5s. During 2- 7s (38- 40 °C), the rate was 2.13 ± 0.48 AP per 5s, giving a p-value of 0.9245; while a rate of 2.53 ± 0.49 AP per 5s was seen during the during 7- 12s (38- 39 °C) duration, with a p-value of 0.6364; and the 17- 22s (37 – 38 °C) period gave a rate of 3.53 ± 0.61 AP per 5s, the p-value being 0.1044. All p-values were calculated by unpaired T-test and n = 15 in all cases (Figure 3G, H).

We quantified the reduction in AP firing in Ano1/TMEM16A^+^, MNP^+^ neurons by calculating the Ca^2+^ activity suppression ratio. To calculate the suppression ratio, we first numerically integrated the bleach corrected, baseline subtracted, normalized GCaMP6f signal. A reduction of the slope of the linear fit of such this integrated GCaMP6f signal curve indicates a decline of Ca^2+^ activities, while an increased slope indicates increasing activity (Figure 4A). The slope of the integrated GCaMP6f signal of Ano1/TMEM16A^+^, MNP^+^ neurons was analyzed during three distinct time periods: one prior to AMF application (baseline Ca^2+^ activity rate at 37 °C, within −20 to −2 s), one during magnetothermal silencing (lying within 2 – 22 s, with temperature > 37 °C) and a final period, when Ca^2+^ activities resumed to baseline levels (between 22 – 30 s). In Figure 4B, a reduced slope is observed in the pooled integration plot, following the AMF application. This corresponds to the silenced period, which is followed by resuming Ca^2+^ activities, revealed by a increasing slope. During the silenced state, the slope was suppressed by 82.04 ± 8.84 % compared to the baseline slope (suppression ratio). This slope during silencing was significantly different from the baseline values (p-value = 0.028 (n = 12, unpaired T-test)). The median suppression ratio was 95.45 %, while the third quartile was at 67.99 %. After normal firing resumed, the slope was indistinguishable from the starting slope (reduction was −8.98 ± 28.98 %) (Figure 4C).

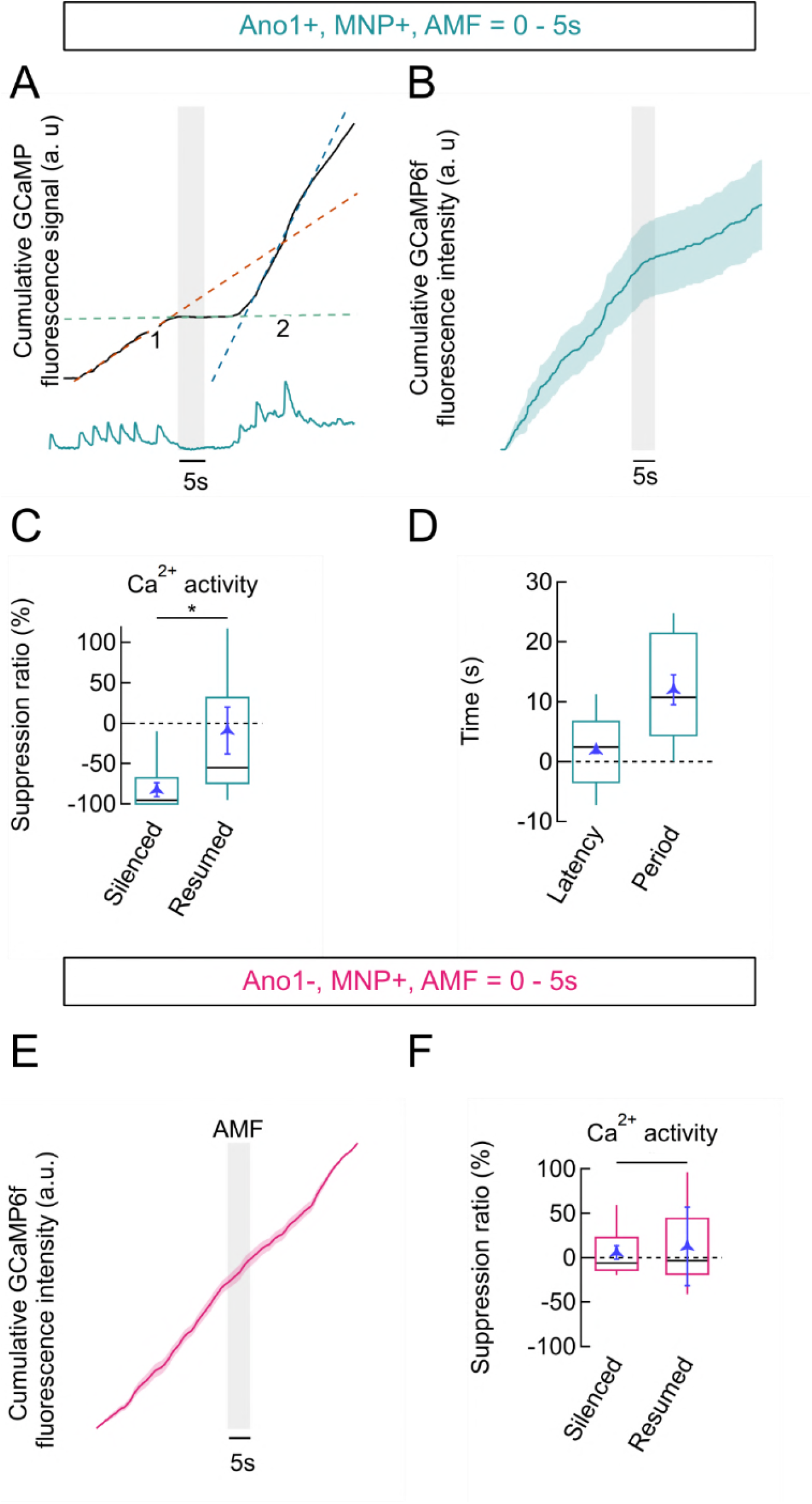
Firing suppression ratio and latency. **(A)** Representative numerical integration (top) of the GCaMP6f plot (bottom), shows suppression with AMF application (grey bar) in a Ano1/TMEM16A+, MNP+ neuron. Suppression is indicated by a reduction in slope of the integration plot. Dotted lines show linear fits of three distinct regions of the trace (red: before AMF, green: during AMF suppression, blue: after resumption, following suppression). Slopes of these lines gives average rate of Ca^2+^ influx, during the indicated periods. The points of intersection of these lines give the times corresponding to the beginning (1) and ending (2) of suppression. **(B)** GCaMP signal integration (trapezoidal) in Ano1/TMEM16A+, MNP+ neurons (mean ± sem, n = 12). AMF is indicated by the grey bar. **(C)** Box plot showing ratios of slopes obtained from integration plots in Ano1/TMEM16A+, MNP+ neurons. Suppression ratio is given as change in slope during suppression (silenced) and after the resumption of firing (resumed), compared to the initial slope. Ratio obtained during the silenced period was −82.037 ± 8.835 % and −8.982 ± 28.977 % during resumption (mean ± sem). The p-value obtained was 0.028, by unpaired T-test. **(D)** Box plot showing the latency of silencing and the period of silencing, following five seconds of AMF in Ano1/TMEM16A+, MNP+ neurons. Latency was 1.882 ± 0.477 s following the start of AMF application, while the period was 12.05 ± 2.477 s (mean ± sem). **(E)** GCaMP signal integration (trapezoidal) in Ano1/TMEM16A^-^, MNP+ neurons (mean ± sem, n = 16). AMF is indicated by the grey bar. **(F)** Box plot showing ratios of slopes obtained from integration plots in Ano1/TMEM16A^-^, MNP+ neurons. The ratios obtained in similar time periods as indicated in (C) yield 6.02 ± 7.74 % and 12.88 ± 44.20 % during the *silenced* and *resumed* periods, respectively. No significant difference between these rates was found (t-test). All AMFs were 30 kA/m r.m.s at 412.5 kHz. Boxes span the second and the third quartile and the whiskers point at 10^th^ and 90^th^ percentile. Black bars in boxes indicate medians.

To determine latency and duration of magnetothermal silencing of Ano1/TMEM16A^+^, MNP^+^ neurons, we determined the time (with AMF start at time = 0s) of the intersection points of the sectional fitting lines (indicated by 1 and 2 in Figure 4A). Intersection of the slope of baseline activity with the slope of the silenced period defines the beginning of inactivation and hence the *latency of si/encing*. Similarly, the intersection of slope during silencing with the slope of the recovered activity defines the end of silencing. We found the latency of silencing in Ano1/TMEM16A^+^, MNP^+^ neurons to be 1.88 ± 0.48 s (after the start of AMF). The end of silencing was observed at 13.93 ± 2.02 s, giving a silencing period of 12.05 ± 2.48 s for 5 s long and 30 kA/m at 412.5 kHz AMF (Figure 4D).

Similar analysis on Ano1/TMEM16A^-^, MNP^+^ neurons (Figure 4E) showed no significant suppression: suppression ratio of 6.02 ± 12.88 % in the period following AMF application (between 2 – 22 s) and 7.74 ± 44.20 % long after the AMF was turned off (between 22 and 30 s), (p-value = 0.615, n = 16, T-test). This ascertains that the silencing observed was unique to Ano1^+^/TMEM16A^+^ neurons.

## Discussion

In this Research Advance, we extended the magnetothermal neuro-modulation technique to silencing in neuronal culture, using the thermosensitive chloride channel Anoctamin1. Similar to neuronal activation, using TRPV1 ion channels, only 2 – 3 degrees above the physiological temperature was sufficient to induce significant suppression in firing. The speed of suppression initiation depends on heating rate of the magnetic nanoparticles attached to the membrane and may be tune by the parameters of the applied alternating magnetic field.

Magnetothermal silencing can easily be extended to in-vivo models using the same approaches as used in our in-vivo demonstration of magnetothermal neuronal activation (Munshi et al, 2017). As Ano1/TMEM16A channels are mammalian ion channels, they target well to the neuronal membrane and are efficient chloride channels (Pedemonte & Galietta, 2014; Terashima et al., 2013). Yet, as we have shown here, overexpressing them in neurons does not alter neuronal activity or shape of calcium transient at normal physiological temperatures. The gene encoding Ano1/TMEM16A is too large to be delivered using AAV, but we have successfully used Lentivirus to deliver Ano1/TMEM16A to neurons (Figure 1 – Figure supplement 1).

Magnetothermal genetic silencing offers a minimally invasive alternative to optogenetic silencing with the advantages of easy deep tissue penetration of AMF, no requirement for any tether or external marking of the animals, and the possibility for prolonged silencing without the undesired effects of blue light induced photo-toxicity (“Artifacts of light,” 2013; Dixit & Cyr, 2003).

## METHODS

### Rat Hippocampal Neuronal Cultures

Primary rat hippocampal neurons were harvested from E18 rat fetuses, from Timed Prenancy female Sprague Dawly rats (Harlan), following a modified standard protocol (Brewer, 1997). All animal protocols were approved by the Animal Care and Use Committee at University at Buffalo, State University of New York. The neuronal cell cultures were used for stimulation experiments between days 8 and 14 in-vitro (DIV).

### Transfection of Hippocampal Neurons

Plasmids encoding Anoctamine1 with an mCherry marker (Ano1/TMEM16A-mCherry) and GCaMP6f were introduced into the hippocampal neurons using Calcium Phosphate method, following published protocols(Jiang & Chen, 2006). Transfection was performed on DIV 4 or 5 using 5 μg Ano1/TMEM16A-mCherry p-DNA and 2 μg GCaMP6f p-DNA in 35 mm dishes (Corning) with four 12 mm poly-L-lysine coated cover slips.

### Preparation of Neurons for Silencing and Imaging

After transfection, the neuronal cultures were placed in incubator (37 °C, 5% CO_2_) for next 48−72 hours before experiment. 24 hours prior to the experiment, 1 μM TTX (Tetrodotoxin, sigma-Aldrich) was added to the culture media. For imaging, the neurons were placed in imaging bath solution (NaCl 145, CaCl2 2, MgCl2 1, KCl 2.5, HEPES 10, Glucose 20 (all in mM) at pH 7.34, and the osmolality was adjusted to 310−315 mOsmole/L.

Synthesized superparamagnetic core-shell Co-Mn-Ferrite nanoparticles (MNP) coated with PMA (poly-isobutylene-maleic anhydride) were functionalized by covalently attaching Neutravidin (Thermo Scientific Cat # 31000) to the PMA layer. Prior to this step, neutravidin was modified to attach dye molecules (Thermo Fisher Cat # A37573). Cells growing on 12 mm cover slips were incubated (10 minutes at 37 °C) with 1.5 μl of biotinylated A2B5 antibody (Invitrogen Cat # 433110) in 200 μl of imaging bath solution. The bath solution was then washed out by perfusion and replaced with 2 μl functionalized nanoparticles in 200 μl of fresh bath solution and incubated for 10 minutes at 37 °C. After this period, the unbound nanoparticles were washed out and imaging was performed, leaving a layer of nanoparticles attached over the entire membrane surface (Figure 2 – Figure supplement 1).

### Live Cell Imaging Under Alternating Magnetic Fields

Alternating magnetic fields (AMF) were generated by a 5 mm Φ, 5 turn water cooled copper coil (Figure 2C) driven by a 7.5 kW ac power supply (MSI Automation). The coil flanked the imaging dish (ALA MS−512DWPW) placed over the microscope (Zeiss Axio Observer A1.0m) objective (Zeiss, working distance, 0.71 mm). Oscillating magnetic fields heat the metallic objective by producing eddy currents, causing focus shift and aberrations. To correct these in the real time, we used a fast, customized piezoelectric autofocus system (Motion X Corporation, 780 nm laser interferometer based) (Figure 2C). A custom-made microenvironment chamber was used to enclose the entire microscope with a stable, externally regulated temperature inside. HBO 200 lamp with appropriate filters were used for illuminating the fluorescent dyes (GCaMP6f: ex, FF02-472/30; DIC, FF495-Di03; em, FF01-525/30. Alexa Fluor 647: ex, FF01- 635/18; DIC, Di02-R635; em, FF01-680/42. All components were Semrock products). Fluorescence data were recorded at 10 Hz, using an Andor NEO sCMOS camera controlled by microManager software (Edelstein, Amodaj, Hoover, Vale, & Stuurman, 2010).

### Data Analysis

The average fluorescence intensity in a ROI (regions of interest) in the fluorescence microscopy data was extracted using FIJI (Fiji Is Just ImageJ). This intensity versus time data was then further processed using IgorPro (WaveMetrics).

The ROI intensity data contained fluorescence signal, offset by dark noise. The dark noise is the mean signal recorded by the camera under no illumination conditions. All experiments were done under similar ambient light conditions and the dark noise value, generally a function of the camera exposure time and binning deviated little from experiment to experiment. A constant dark noise value was subtracted from all data points. Bleach correction was subsequently done, if the baseline showed substantial bleaching based decay. For this, the baseline was fitted to an exponential function and a modified signal was obtained according to Equation (2).

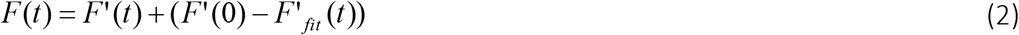

Here, *F’(t)* is the signal values obtained after dark noise cancellation and *F’_fit_ (t*)) is the corresponding exponential fit function. Data, thus modified contained calcium peaks over a constant baseline. The data was then normalized and converted to percentage change in fluorescence. The percentage change in fluorescence intensity was given by Equation (3).

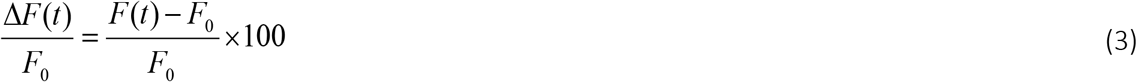

where, *Fo* is the bleach corrected baseline ROI intensity and *F(t)*, the intensity at any time *t*.

### Reconstructing the Firing Pattern from GCaMP6f Calcium Signals

The signal gain and temporal resolution of GCaMP6f makes it sensitive enough for single action potential detection. However due to the long decay time of GCaMP6f, it is challenging to isolate AP spikes from an AP train (Chen et al., 2013). To extract action potential events from a GCaMP6f fluorescence signal (Ca^2+^ peaks), we first isolated all single peaks from the intensity normalized (and baseline corrected) data. Isolated spikes produce a GCaMP6f peak intensity of about 5% of the baseline. Pooling such isolated spike related GCaMP traces, we obtained the average GCaMP peak, corresponding to single AP firing (Figure 1 – figure supplement 2B). The average spike wave was then interpolated linearly to increase the temporal resolution to 100 Hz.

A null wave of time length equal to that of the original data, but of 100 Hz frequency (in contrast to 10 Hz, in the original data) was created. The null wave was then converted to a binary wave, with ones at the location of estimated AP spikes (Figure 3 – figure supplement 1A). The estimated calcium profile from the single spike (from the previous section) was then convolved with the binary wave. An overlay of the normalized data and its corresponding convolution based reconstruction is shown in Figure 3 – figure supplement 1A. A histogram of the residuals of the original and the reconstructed waves was fitted to a Gaussian function (Figure 3 – figure supplement 1B). The locations of the ones in the binary wave were adjusted to obtain an overall position error (σ) of less than 2%. The binary wave, obtained after optimization gave the temporal function of AP events for each recording.

Average firing rates (in Hertz) were found in two ways. The total number of events in a given time interval were counted. Dividing the total events by the time interval gave the frequency of AP firing in Hertz (Figure 3). Another estimation of firing rate was done by first numerically integrating (trapezoidal) the spike train data (binary wave). Integrated values were obtained at each point of the integrand; giving the integrated wave a constant time scaling of 0.01 s. Linear fit of the integration trace over fixed regions, corresponding to the time intervals of interest was then done (Figure 4). The slope of the line thus obtained, gave the probability of finding an action potential in the given time interval. Since, data points were 0.01 s apart, the average frequency of AP firing (in Hz) over the time span was given by

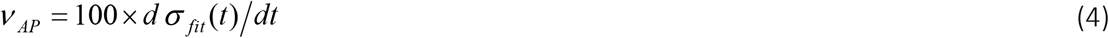

Here, ^σ_fit_(t)^ is the coefficient of the first order term of the linear fit of the integrated spike train wave at any time *^t^*, and *^V^AP* is the average frequency of spiking in within the time interval.

### 2.6.1 Finding the latency and duration of silencing

To find the silencing latency and duration, individual spike train integration plots were divided into three temporal sections, for linear fitting. The first section represented the state before the AMF was applied. The second section,covered the period of inactivation (silencing), while the third section represented the period when firing had resumed after inactivation. The point of intersection of the fitting lines of the first and the second sections gave the starting time of activation, while the intersection of the fitting lines of the second and third sections gave the ending time of the inactivation period. Average start and end times were calculated from the times obtained from individual plots(Figure 4D).

## Acknowledgements

We thank Sara D. Parker and Jason Myers (University at Buffalo) for molecular cloning and culturing of hippocampal rat neurons. pGP-CMV-GCaMP6f was a gift from Douglas Kim (Addgene plasmid # 40755); mANO1/TMEM16A-eGFP and mANO1/TMEM16A-mCherry were gifts from H. Criss Hartzell, Jr. (Emory University). This work was supported by NIH grants was supported by HSFP project grant RGP0052/2012 and NIMH grants R01-MH094730 and R01-MH111872.

## Author contributions statement

The project was conceived by AP. AP and RM authors contributed to design of the research. RM performed cell silencing experiments and the imaging. RM analyzed GCaMP6f data, wrote the code to do so, and prepared figures. ICR prepared the MNPs. RM and AP wrote the manuscript. All authors reviewed the manuscript.

## Additional information

The authors declare no competing financial interests.

---

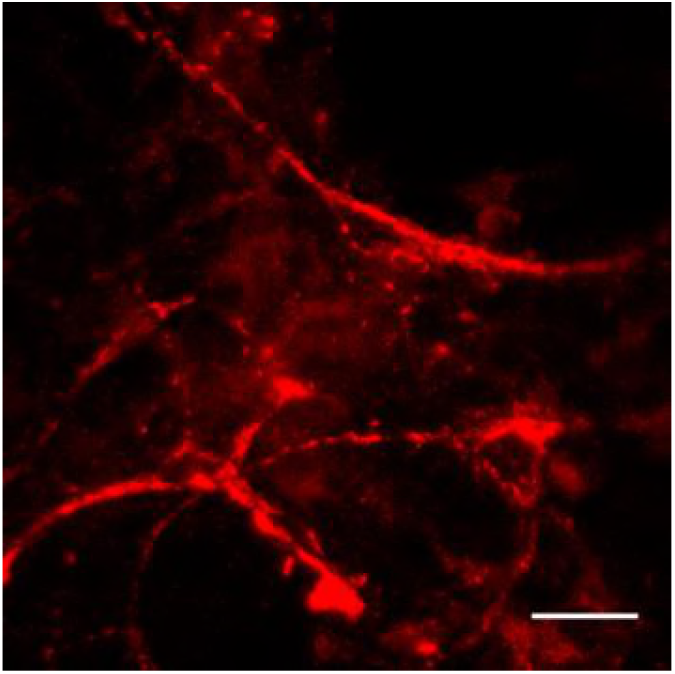
Ano1/TMEM16A-mCherry expression 72 hours after incubation in neurons infected with ano-mcherry lentivirus.

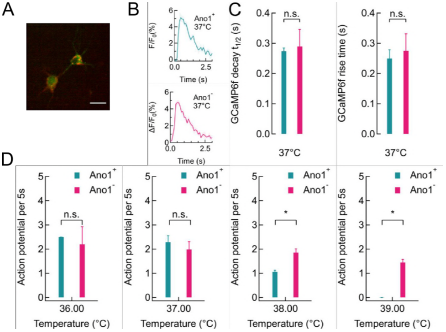
**(A)** Hippocampal neurons cotransfected with ano-mCherry and GCaMP6f DNA plasmids. **(B)** Average single gCAMP6f peaks recorded in Ano1 ^+/-^ neurons (A and B respectively) at 37 °C respectively (n = 4). **(C)** No significant change in peak characteristics was found (n = 4, all cases). **(D)** Firing rates of Ano1 ^+^ and Ano1^-^ neurons were not significantly different at 36 (p = 0.6628) or 37 °C (p = 0.4526), while firing rates at 38 and 39 °C varied significantly between Ano1^+^ and Ano1^-^ neurons (p = 0.0227 and 0.0051 respectively).

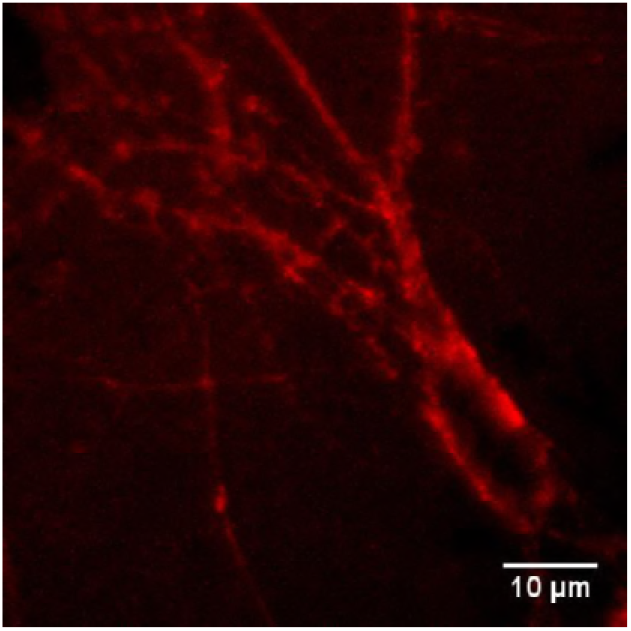
Fluorescence micrograph showing magnetic nanoparticles coated with Alexafluor-647 labeled NeutraAvidin attached to the neuronal membrane in a rat hippocampal culture via biotinylated anti-A2B5 antibody

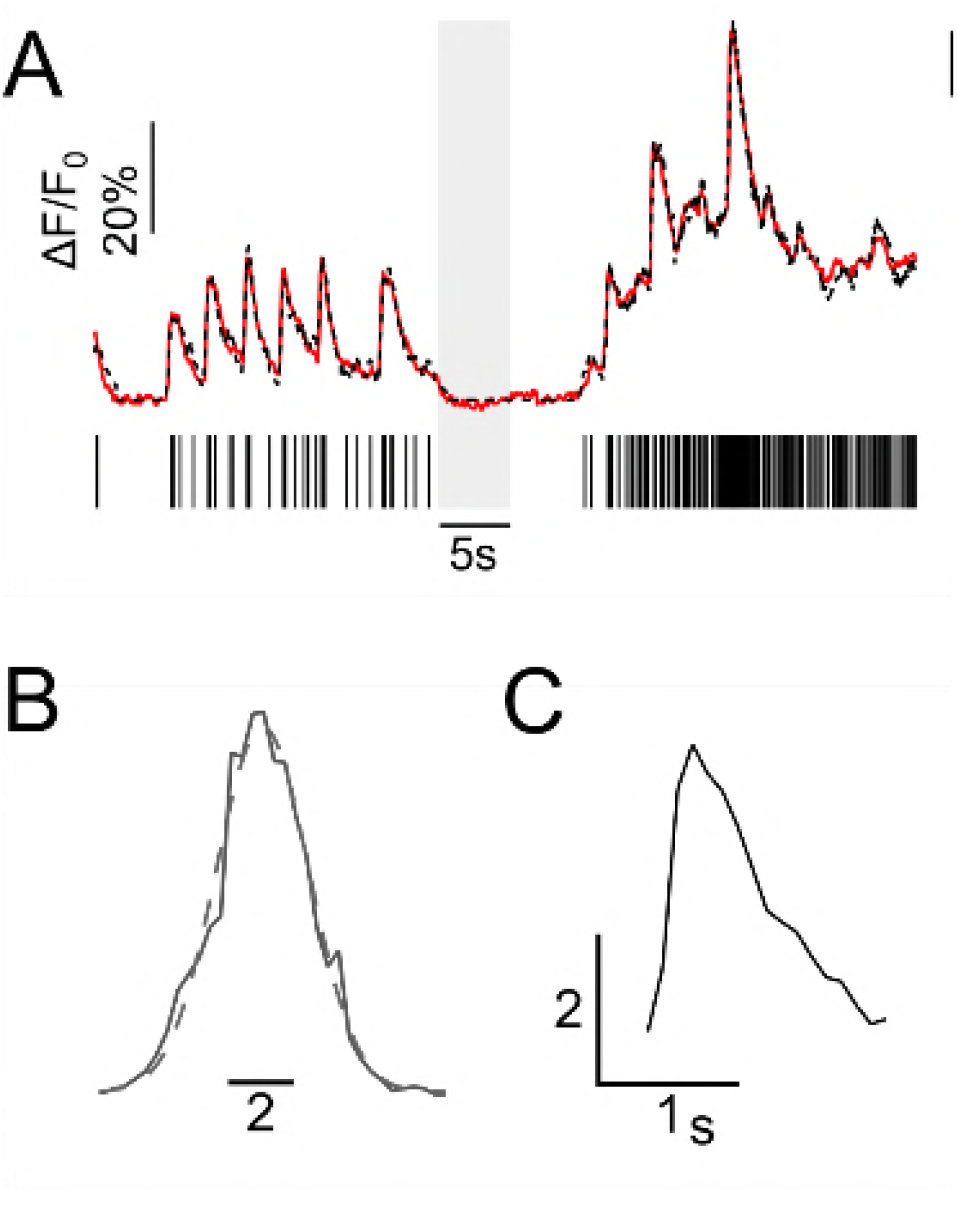
**(A)** Predicted action potential events from GCaMP6f trace (black sticks) and the overlay of regenerated trace obtained from convolving predicted action potential events with single AP calcium peak (black, broken) with the normalized GCaMP6f trace. **(B)** Histogram of the residual of the GCaMP6f trace and the regenerated trace in (A), fitted with a Gaussian curve. The sigma of the fit was 1.42 ± 0.05. **(C)** The average (n= 3 peaks) signal from three smallest peaks recorded from the same neuron.

